# Fast-Part: Fast and Accurate Data Partitioning for Biological Sequence Analysis

**DOI:** 10.1101/2024.11.13.623463

**Authors:** Shafayat Ahmed, Muhit Islam Emon, Nazifa Ahmed Moumi, Liqing Zhang

## Abstract

Developing effective machine learning models for classifications of biological sequences depends heavily on the quality of the training and test datasets split. Existing tools are either computationally expensive, unable to maintain the desired level of similarity between the training and test datasets, or unable to retain training-test ratio stratification. Here, we present Fast-Part, a fast and accurate sequence data partitioning tool that ensures strict homology separation between the training and test datasets and the best possible training: test stratification ratio, and at the same time, is computationally fast. Fast-Part demonstrates rapid and accurate partitioning performance across diverse protein sequence datasets and maintains strict partitioning compared to the existing tools. Fast-Part can handle massive datasets and maintain strict homology partitioning.

## 1 Introduction

Partitioning of biological sequences into training and testing sets is crucial in developing robust machine-learning models for sequence classification. Random splitting can lead to the overestimation or underestimation of model performance. Homology between sequences is quantified using DIAMOND sequence alignments with user-specified identity threshold and query coverage.Sequences in the train and test sets are considered homologous and flagged if their alignment identity exceeds the threshold and coverage exceeds the specified value. While evaluating it’s paramount making sure all cross-set alignments fall below the defined similarity boundary. Among common practices, one of them is stratified k-fold cross-validation. In this approach, the dataset is split into k partitions where sequences are assigned to each partition randomly, and in turn, each of these partitions serves as the test set, with the remaining partitions acting as the training set. The overall accuracy is the mean accuracy across all k-folds[15]. Sometimes, instead of multiple partitions, datasets are split into two (train and test) or three sets (train, test, and validation), with the train set typically comprising 60-80% of the data. These methods often fail to maintain sequence homology between the train and test sets, potentially overestimating the model’s accuracy[14].

An alternative partition method is based on sequence homology and involves clustering the dataset such that each cluster maintains up to a pre-defined similarity threshold with the other clusters. If the threshold is set at 60%, no sequences from different clusters should have a similarity of more than 60%. Popular tools such as CD-HIT[3] and MMseqs2[13] use this approach and can cluster thousands of sequences in a short period. In particular, CD-HIT calculates sequence similarity by counting shared k-mers and grouping sequences into clusters based on a specified similarity threshold, with the longest sequence initially chosen as the reference. For sequence similarities below 40%, CD-HIT switches to PSI-CD-HIT mode, which utilizes PSI-BLAST for more refined clustering analysis, as described by Hobohm et al.[4]. MMseqs2[13] uses a substitution matrix to compare k-mers and identify high-scoring pairs aligned locally, similar to BLASTP’s methodology. Its default clustering mechanism, a greedy set-cover algorithm, selects sequences that connect with most others in the dataset, though it can also employ CD-HIT’s strategy for grouping. Both methods group similar sequences into clusters, each with a lead sequence representing all the sequences within that cluster. This reduces computational costs because only the lead sequence is processed instead of every sequence in the cluster. However, this approach does not guarantee that sequences from different clusters have a consistent level of similarity to each other; that is, it ensures similarity within clusters but not necessarily between clusters. GraphPart[14] addresses this issue by proposing a method that compares pairwise sequence similarity and clusters sequences with a strict similarity threshold identity. While earlier methods for reducing sequence homology relied on global[8] or local[12] alignment techniques using metrics like percentage identity and overlap length to gauge similarity[11, 7, 10] GraphPart[14] advances this approach by comparing all pairwise sequence similarities and enforcing a strict similarity threshold during clustering. This method, inspired by Hobohm and colleagues’ algorithm from 1992[4], requires calculating all pairwise sequence similarities, which is computationally expensive and makes clustering large datasets challenging. Moreover, maintaining a strict similarity threshold across the entire dataset often fails to preserve the stratification ratio for each class. For instance, in classification tasks, significant deviations from the intended train-test ratio for certain courses can lead to underperformance due to insufficient training data or an excessively large test dataset, resulting in skewed model evaluations [8, 12, 7, 10]. Other recent tools like DataSAIL [5] preserve sequence similarity and maintain class stratification across splits, enabling robust evaluation on heterogeneous data but adding complexity for homogeneous datasets.

Additionally, challenges such as class imbalance and the presence of many classes with small sample sizes demand careful consideration. An effective and fair split of the dataset, taking into account these challenges, is crucial for maintaining the quality of model training and validation processes. We addressed these issues with a fast and accurate data partitioning tool that maintains: (1) a strict similarity threshold between sequences from the same class label while ensuring no sequence in the test set has similarity with train sequences above the threshold, (2) time efficiency, comparable to CD-HIT/MMseqs2, capable of handling large data, (3) close adherence to the stratification ratio for each class., with a minimal margin of error, (4) retention of data as much as possible through iterative reassignment, and (5) a randomized but balanced distribution to ensure the training data encompasses a wide variety and captures more patterns. Our model demonstrated superior performance over other partitioning tools in achieving these objectives.

## 2 Materials and Methods

### 2.1 Datasets

We benchmarked our sequence partitioning tool using four diverse protein sequence datasets, as shown in Table 1. These datasets include the antimicrobial resistance gene classification dataset HMD-ARGDB[6], the subcellular localization dataset DeepLoc[1] that includes eukaryotic subcellular localization classification of proteins, a Metal Resistance Gene classification dataset[9], and the large-scale mobileOG-db dataset[2] of mobile orthologous groups.

**Table 1:** Total number of sequences, number of labels, max and min length of the sequences for the four datasets.

| Dataset | Number of Labels | Size | Max. Length | Min. Length |
| --- | --- | --- | --- | --- |
| HMD-ARGDB | 33 | 18k | 1576 | 34 |
| METAL | 1 | 155k | 1895 | 47 |
| Mobile-OGDB | 6 | 1M | 14944 | 29 |
| DeepLoc | 25 (7) | 14004 | 3579 | 69 |

We selected these four representative and diverse biological sequence datasets to evaluate Fast-Part under a wide range of real-world conditions:

- HMD-ARGDB (Antimicrobial Resistance Genes): Chosen for its multi-class, imbalanced nature (33 labels, some with very few sequences). This dataset poses a significant challenge for homology-based partitioning, making it ideal to evaluate stratification and homology enforcement under imbalance.
- DeepLoc: A well-curated dataset for eukaryotic subcellular localization, containing 25 classes and moderately sized sequences. It allows us to test how well Fast-Part performs on relatively balanced, moderately complex datasets used in protein localization tasks.
- Metal Resistance Gene (MRG) dataset: This single-label dataset helps evaluate Fast-Part’s efficiency and correctness in simple scenarios where stratification is trivial but homology separation must still be enforced.
- mobileOG-db: With over 1 million sequences and six labels, this large-scale dataset challenges the scalability and runtime performance of partitioning tools. It is particularly useful for benchmarking computational efficiency.

This dataset selection ensures evaluation across - scale (from 14k to 1 million sequences), label complexity (single vs. multi-class), length distribution (short to long proteins), homology structure (dense to sparse sequence similarities).

### 2.2 Fast-Part Architecture

Figure 2, presents an overview of the Fast-Part data partitioning tool, designed to meet the specified objectives. The tool accepts a FASTA file, along with parameters such as a similarity threshold, coverage, and a desired stratification ratio as inputs. The output consists of train and test FASTA files, strictly maintaining a similarity threshold between the sets. The tool also ensures that the stratification ratio for each label closely aligns with the user-specified ratio. Additionally, Fast-Part generates a list of sequences excluded during processing and a text file containing the names of labels that could not be partitioned within the established thresholds for similarity and stratification.

**Fig. 1:**
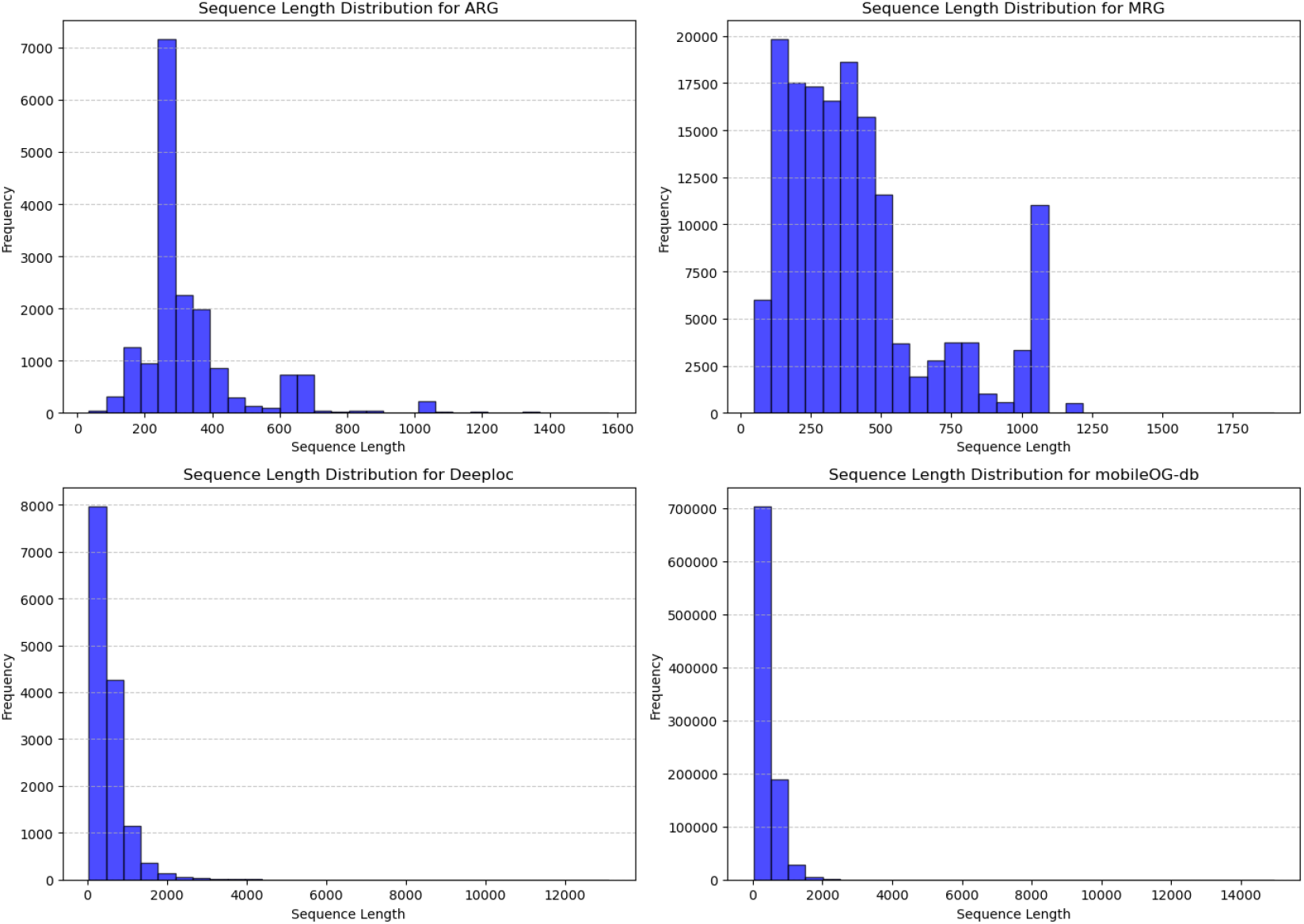
Sequence length distribution for the four datasets: ARG, MRG, Deeploc, and mobileOG-db.

**Fig. 2:**
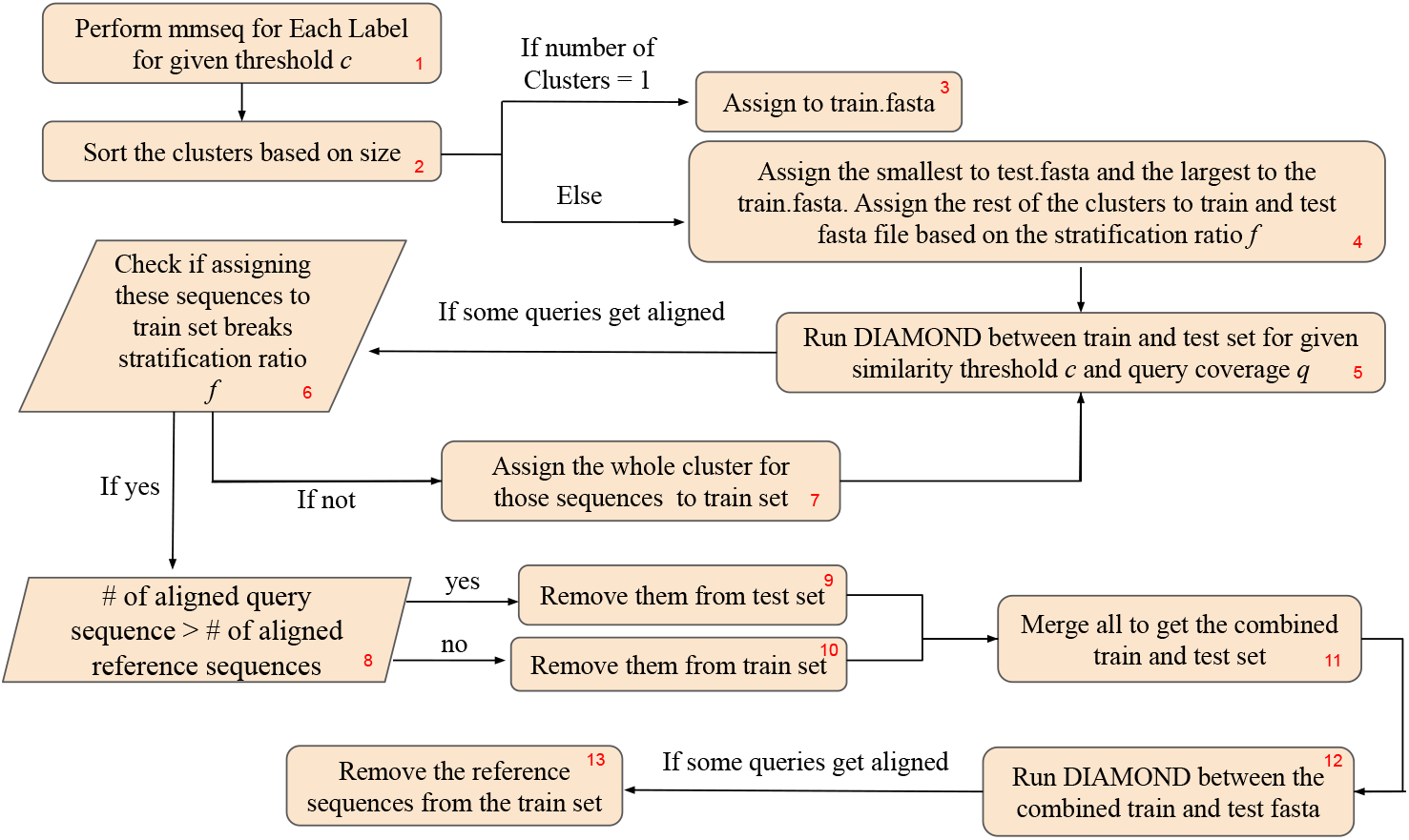
A Flowchart of the Fast-Part data partitioning pipeline.

Algorithm 1, outlines our pipeline, which is structured into four critical steps, each contributing to the robustness and efficiency of the partitioning process. Each step will be detailed with the methodologies employed.

#### 2.2.1 Division

The initial step involves processing the provided FASTA file and splitting it into one or more FASTA files, each containing only the sequences associated with a single label. (Figure 2(1), Algorithm 1(line 1,2))

#### 2.2.2 Clustering

The next step clusters each FASTA file using CD-HIT or MMseqs2, depending on the selected similarity threshold. Fast-Part uses MMseqs2 to speed up clustering. Each FASTA file produces one or more clusters. If only one cluster is found, it means the file cannot be split at the given similarity threshold, so this cluster is assigned to the training set. If multiple clusters are identified, the largest cluster and its representative sequences go to the training set, while the smallest is assigned to the test set. This decision is particularly important for label groups with few sequences, where random assignment might inadvertently place the largest cluster in the test set, leaving the training set with insufficient examples to learn from. By assigning the largest cluster to training, we ensure that the model is trained on the most representative portion of such underrepresented labels. Moreover, clusters are formed based on high sequence similarity, meaning that sequences within a cluster are often highly redundant. Whether the larger or smaller cluster is used for training or testing matters less in such cases, as the internal diversity is low. Therefore, assigning the largest cluster to the training set does not introduce significant bias.

##### Algorithm 1

Fast-Part Sequence Clustering

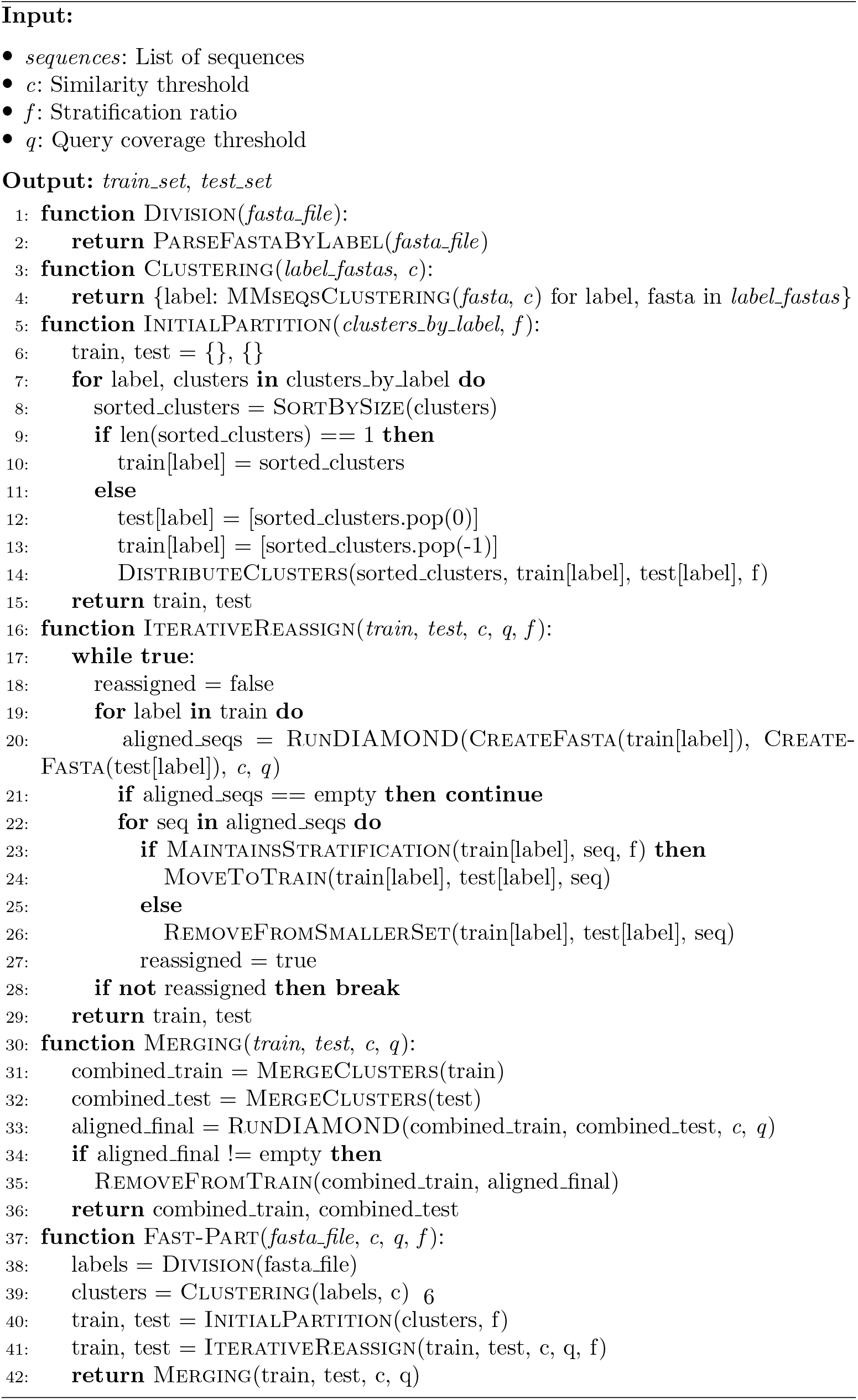

Remaining clusters are distributed between the training and test sets based on the specified stratification ratio. Since CD-HIT and MMseqs2 prioritize high in-cluster similarity, we initially assign clusters slightly below the target ratio to allow finer control during final sequence assignment. (Figure 2(1-4), Algorithm 1(line 3-15))

#### 2.2.3 Iterative Reassignment

Following the initial partitioning based on similarity thresholds and stratification ratios, we assess the partitions for sequence similarity using DIAMOND alignment between the train and test FASTA files for each label. If the DIAMOND output indicates similarities above the similarity threshold between train and test sequences, we flag those sequences in the test set and their respective clusters for potential reassignment. We then evaluate whether transferring these sequences to the training set would alter the stratification ratio for that label. If such a transfer changes the ratio, we remove the implicated sequences, choosing those from the test set if they are fewer than their counterparts in the training set, and vice versa. If the stratification ratio is unaffected, the sequences are reassigned to the training set. This process is repeated until no sequences in the test set align with those in the training set (i.e., having similarity above the threshold). (Figure 2(5-10), Algorithm 1(line 16 - 29))

#### 2.2.4 Merging

Upon completion of the iterative reassignment, we compile the train and test files for each label into comprehensive train and test sets. These consolidated sets are structured to prevent any similarity above the defined threshold between sequences of the same label. However, sequences from different labels having homology above the threshold may still be present. To address this, a final DIAMOND alignment is conducted between the combined train and test sets using the established similarity threshold and coverage criteria. Sequences from the training set that align with those in the test set are then removed.

This process ensures that dataset partitioning respects both the similarity threshold and the stratification ratio while remaining computationally efficient. It also preserves the integrity of the dataset, enabling reliable and unbiased evaluation of machine learning models. (Figure 2(11-13), Algorithm 1(line 30-36))

## 3 Evaluation

### 3.1 Performance Metrics

Quantitative metrics are extensively reported based on runtime, number of sequences retained, stratification balance [Table 2] and sequence homology.

**Table 2:** Label stratification categories based on train-test partition characteristics.

| Category | Description |
| --- | --- |
| **Balanced** | Train-test split deviates less than or equal to 10% from the desired stratification ratio. |
| **Moderate** | Deviation between 10% and 20% from the desired stratification ratio. |
| **Low Sample** | Labels with fewer than 15 sequences. A 50-50 split is marked as Balanced; otherwise categorized here. |
| **Ambiguous** | Only train or test set is present. Occurs if all sequences fall into a single cluster or are removed due to high similarity with other label samples. |
| **Other** | Poorly stratified labels with more than 20% deviation from the target train-test ratio and not fitting into above categories. |

To maintain consistency across datasets all sequences were converted to standard FASTA format. Sequences with ambiguous or invalid amino acids were filtered out. Redundant sequences within a label were removed using CD-HIT at 100% identity to avoid skew in cluster sizes. For benchmarking, datasets were subjected to uniform similarity thresholds (e.g., 30%, 40%, 60%, 80%) and a target stratification ratio of 80:20 train-test split unless otherwise specified. These steps ensured fair and comparable evaluation across datasets of varying complexity.

To evaluate the performance of Fast-Part, we tested it on a diverse set of datasets. These included datasets with many classification labels, few labels, and varying sequence counts, from small to large. We also analyzed runtime and measured how different similarity thresholds and stratification ratios affected the partitioning outcome across all datasets.

Additionally, we benchmarked Fast-Part against leading data partition tools such as GraphPart[14]. Our comparative studies demonstrated Fast-Part’s superior performance, establishing its effectiveness in data partitioning tasks across diverse conditions.

Fast-Part allows users to define a similarity threshold, which determines the maximum allowable sequence identity between the training and test sets. In our study, we selected four thresholds 30%, 40%, 60%, and 80% to reflect a representative range of practical use cases in biological sequence analysis:

30% (Very strict): Minimizes redundancy and potential information leakage. This setting ensures that test sequences are highly dissimilar from the training data, which is useful for evaluating generalization on remote homologs. However, it may result in excluding a larger number of sequences, especially in conserved gene families.

40–60% (Moderate): These thresholds offer a balanced approach commonly used in tasks such as antimicrobial resistance gene (ARG) classification and ortholog detection. They maintain biological relevance while still reducing redundancy across partitions.

80% (Relaxed): Allows closely related variants to be split between training and test sets. This is suitable when biological redundancy is acceptable or when the class boundaries are inherently broad.

By benchmarking Fast-Part across these diverse thresholds, we simulate real-world scenarios that researchers often face. Varying the thresholds enables a more comprehensive evaluation of model performance from stringent to lenient partitions, providing insight into how predictive accuracy changes under different homology constraints. Importantly, this approach offers practical benefits: for any new query sequence, researchers can compute its similarity to the training data and, based on the corresponding threshold, estimate the expected model performance. This enables more informed interpretation of predictions and model confidence. Evaluating a model only under a very strict threshold (e.g., 30%) may underestimate its general performance, while a relaxed threshold may overestimate it. Hence, it is essential to benchmark models across a spectrum of similarity constraints. Fast-Part is designed to make this process efficient, standardized, and biologically meaningful.

## 4 Results and Discussion

At first, we experimented with the four datasets for different similarity thresholds and conducted a detailed analysis of the train-test split’s performance metrics.

For analyzing the stratification ratio of all the labels of each dataset, we categorize them into five different categories -

### 4.1 ARG Dataset

The ARG dataset downloaded from HMD-ARGDB has 33 different classes. We performed a 40%, 60%, and 80% similarity thresholding on this dataset. The dataset is highly imbalanced as 19 classes have altogether only 214 sequences, making it difficult to split into train and test sets while maintaining the thresholds and stratification ratio. Figure 3 shows the performance of Fast-Part for the most challenging dataset among the four ARG datasets for a 40% similarity threshold. Despite these challenges, Fast-Part achieved a 53.1% balanced distribution as well as a 25% moderate split for the dataset. 18.8% (five of the labels) have either missing train or test samples, as it was impossible to split them for a 40% similarity threshold. All of these labels had a small number of samples.

**Fig. 3:**
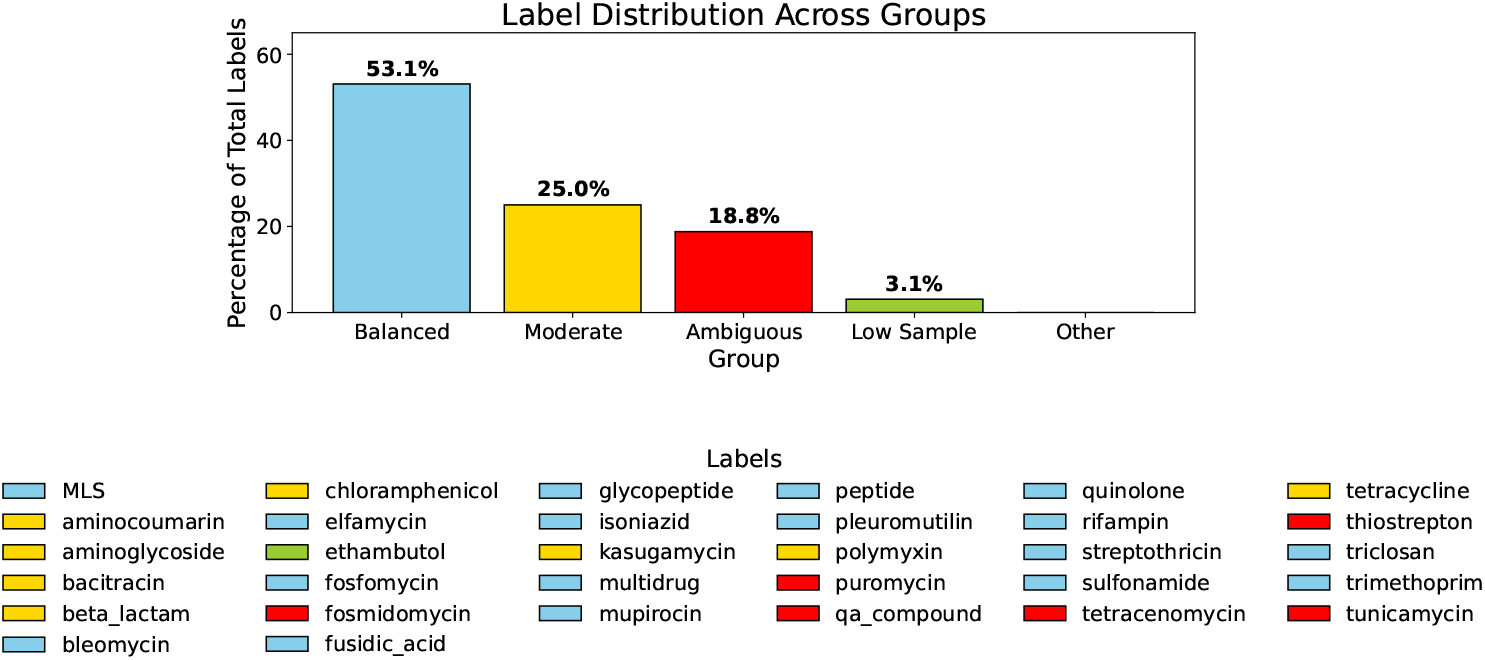
Stratification ratio for the ARG labels for 40% similarity threshold.

As noted, the HMD-ARGDB dataset exhibits significant class imbalance, with 19 of the 33 labels containing fewer than 15 sequences. Under a stringent similarity threshold such as 40%, maintaining both label-wise stratification and homology separation becomes increasingly difficult. This results in several classes falling into one of two scenarios:

- **Single Cluster Classes:** All sequences of a label cluster together due to high similarity, preventing partitioning.
- **High Cross-Homology:** DIAMOND alignment reveals cross-set similarities that cannot be resolved without eliminating the entire class from either the train or test set.

These situations result in labels being marked as ambiguous in our stratification analysis (Figure 3), meaning they lack either a train or test representation after partitioning.

For downstream model training and testing, we exclude ambiguous classes where no valid split is achievable. Including such classes with no representation in one set would lead to biased learning or evaluation failure. The number and identity of excluded classes are recorded in the output log file for transparency and reproducibility.

For 60% and 80% similarity thresholds the model performed well considering there were 33 labels in the ARG dataset. Figure 4 and 5 shows the performance of Fast-Part with 60% and 80% thresholds. Fast-Part achieves 72.7% and 90.9% balanced distribution of the labels for the 60% and 80% thresholds, respectively.

**Fig. 4:**
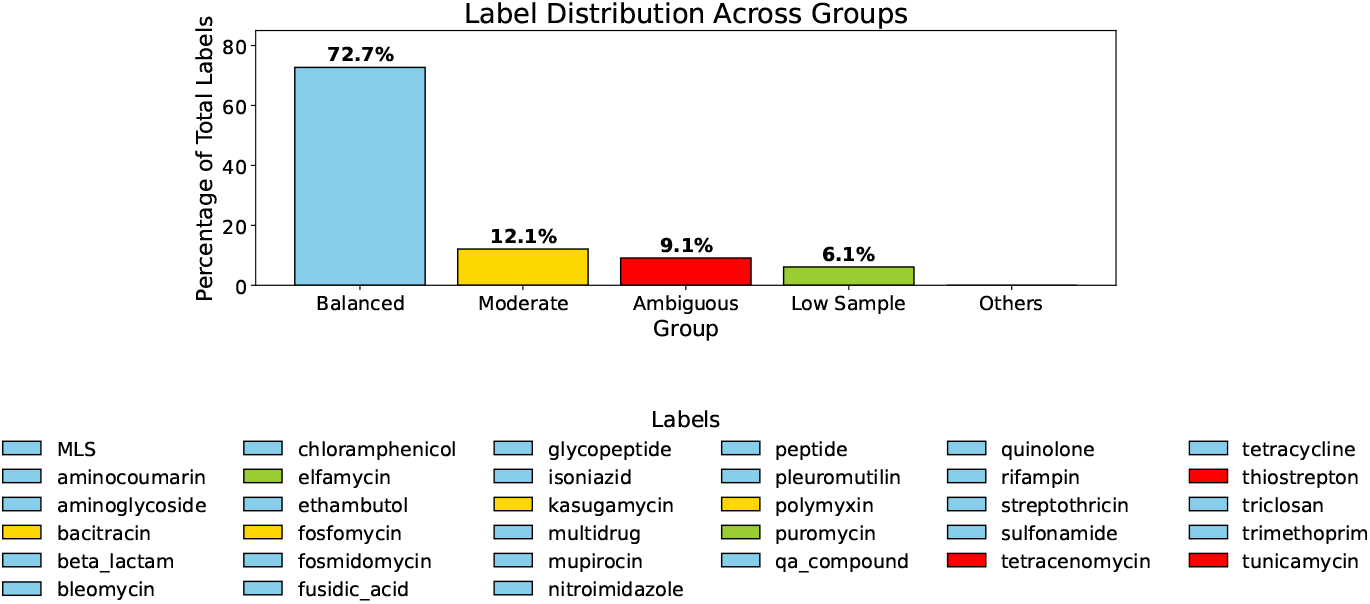
Stratification report for the ARG dataset at 60% similarity threshold.

**Fig. 5:**
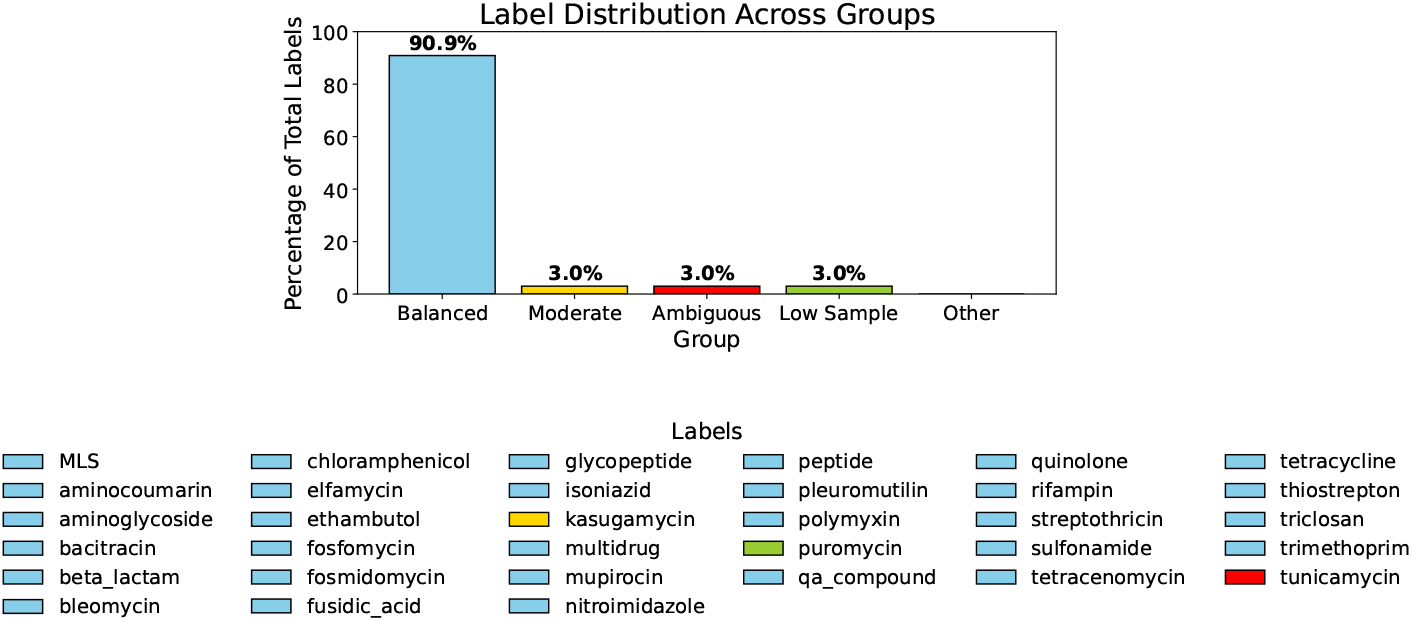
Stratification report for the ARG dataset at 80% similarity threshold.

### 4.2 Deeploc

Fast-Part demonstrated consistent performance on the DeepLoc dataset across various similarity thresholds, maintaining a balanced 80-20 train-test split for all labels except Golgi-apparatus-S (see Supplementary Figure 1).

### 4.3 mobileOG-db

Fast-Part achieved consistent performance in partitioning the large mobilOG-db dataset, which includes over a million sequences across six labels. Despite the dataset’s size, Fast-Part successfully maintained the stratification ratio, achieving balanced splits for all labels at similarity thresholds of 40%, 60%, and 80% [Supplementary Figure 2].

### 4.4 MRG

The final dataset, the metal resistance gene dataset, contains only one label. Compared to the other three datasets, this one presented the least challenge, and Fast-Part achieved a near-perfect split (Figure 6), very close to 80-20, at all similarity thresholds (40%, 60%, and 80%).

**Fig. 6:**
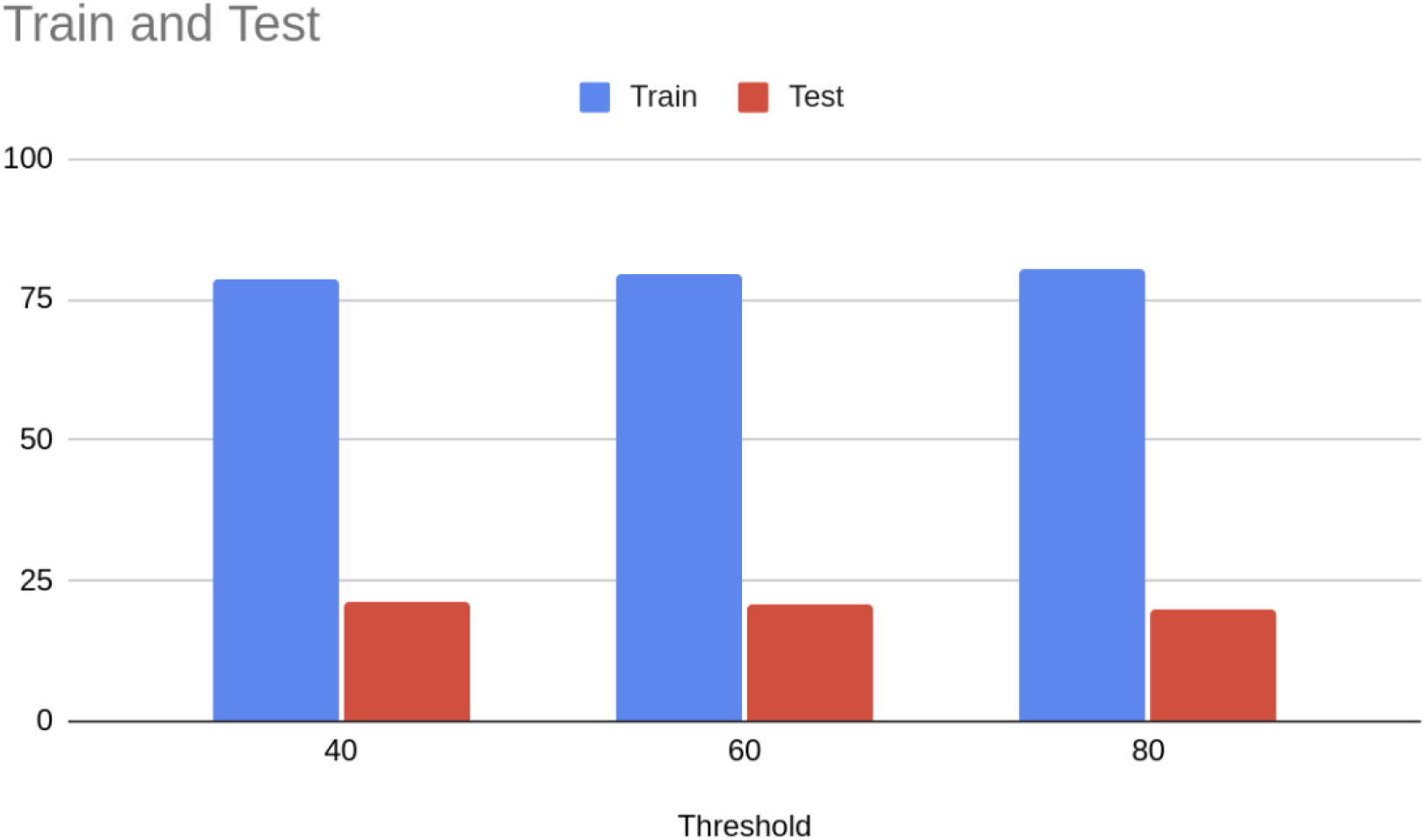
Stratification report for the MRG dataset at 40,60, and 80 percent similarity threshold.

### 4.5 Sequence Retention Analysis

Fast-Part was designed to preserve as many sequences as possible. However, due to the inherent characteristics of the datasets and the requirement to adhere to strict similarity thresholds, Fast-Part had to discard some sequences to ensure balanced distributions. For the deeploc dataset, fewer than 3% of the sequences were omitted even at the 40% similarity threshold. For the metal resistance genes (MRGs), the removal was less than 10% across all thresholds. For the mobilOG-db dataset, about 13% of the sequences were eliminated at the 40% threshold, and the percentage of removed sequences was reduced to less than 2% at the 60% threshold and close to zero at the 80% threshold. For ARGs, the removal rate was consistently less than 15% at all thresholds, a notable achievement considering the challenge of maintaining both stratification ratios and similarity thresholds (Table 4).

### 4.6 Comparison with Other Tools

We compared the efficiency of Fast-Part with that of CDHIT, MMseqs2, and Graph-Part. Three datasets - ARGs, Deeploc, and the NetGPI dataset were used for data partitioning at a strictly 30% similarity threshold for comparison.

GraphPart has two different modes - needle and MMseqs2. The needle mode is recommended but has quadratic time complexity, whereas MMseqs2 is faster but less accurate.

GraphPart and Fast-Part achieve a completely balanced distribution for the Net-GPI dataset. For the Deeploc dataset, GraphPart with the needle mode and Fast-Part achieve complete balance, whereas GraphPart with the MMseqs2 mode achieved 80% balance distribution (Figure 7). However, for both of these datasets, Fast-Part was at least 2x faster than GraphPart with the needle mode (Table 3). We further tested these models on the challenging ARG dataset and the large mobileOG-db dataset. Graph-Part with the needle mode achieved only an 18% balanced distribution and with the MMseqs2 mode got an overall 33% balanced distribution for the ARG dataset (Figure 8). In contrast, Fast-Part got a 51% balanced distribution (Figure 9).

**Table 3:**
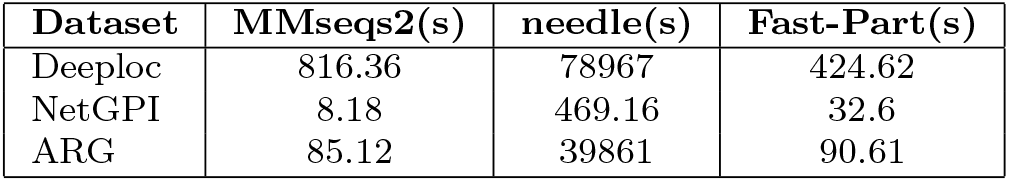
Time taken for splitting into train and test set for the 2 GraphPart models and Fast-Part for the three datasets in seconds.

| Dataset | MMseqs2(s) | needle(s) | Fast-Part(s) |
| --- | --- | --- | --- |
| Deeploc | 816.36 | 78967 | 424.62 |
| NetGPI | 8.18 | 469.16 | 32.6 |
| ARG | 85.12 | 39861 | 90.61 |

**Fig. 7:**
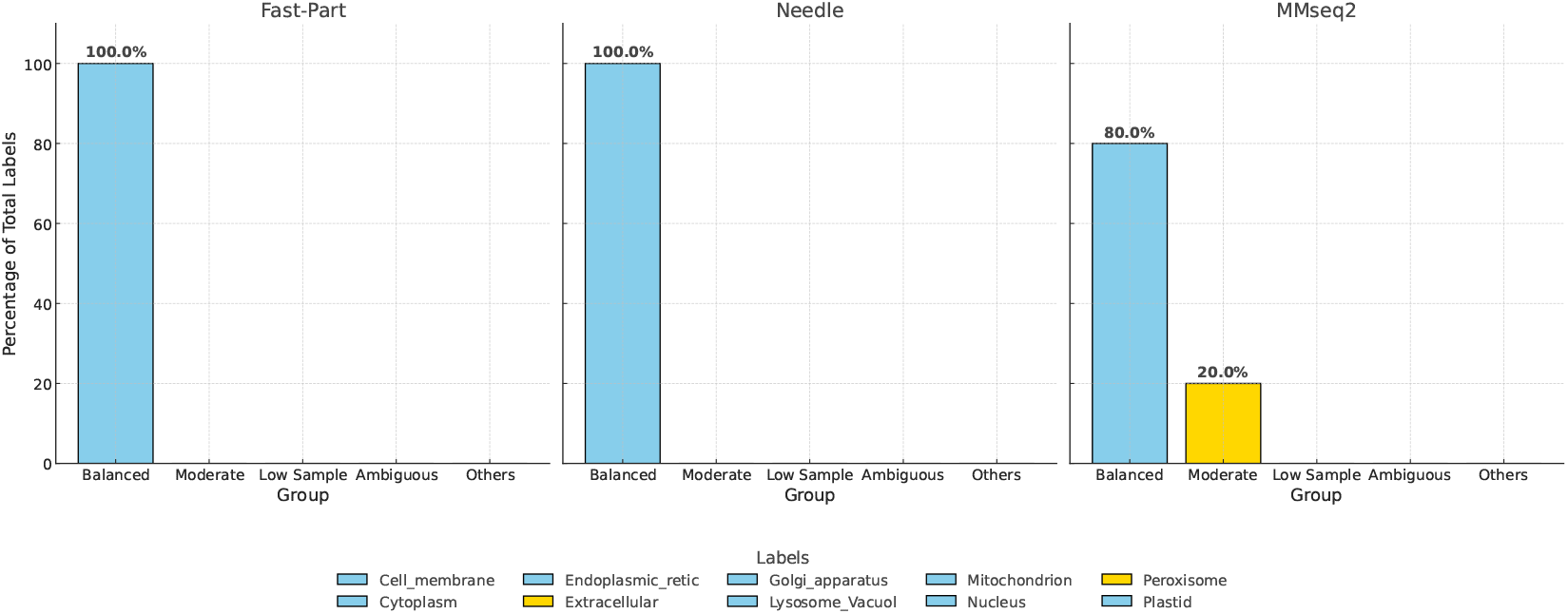
GraphPart and Fast-Part Accuracy on 30% similarity threshold for the Deeploc Dataset.

**Fig. 8:**
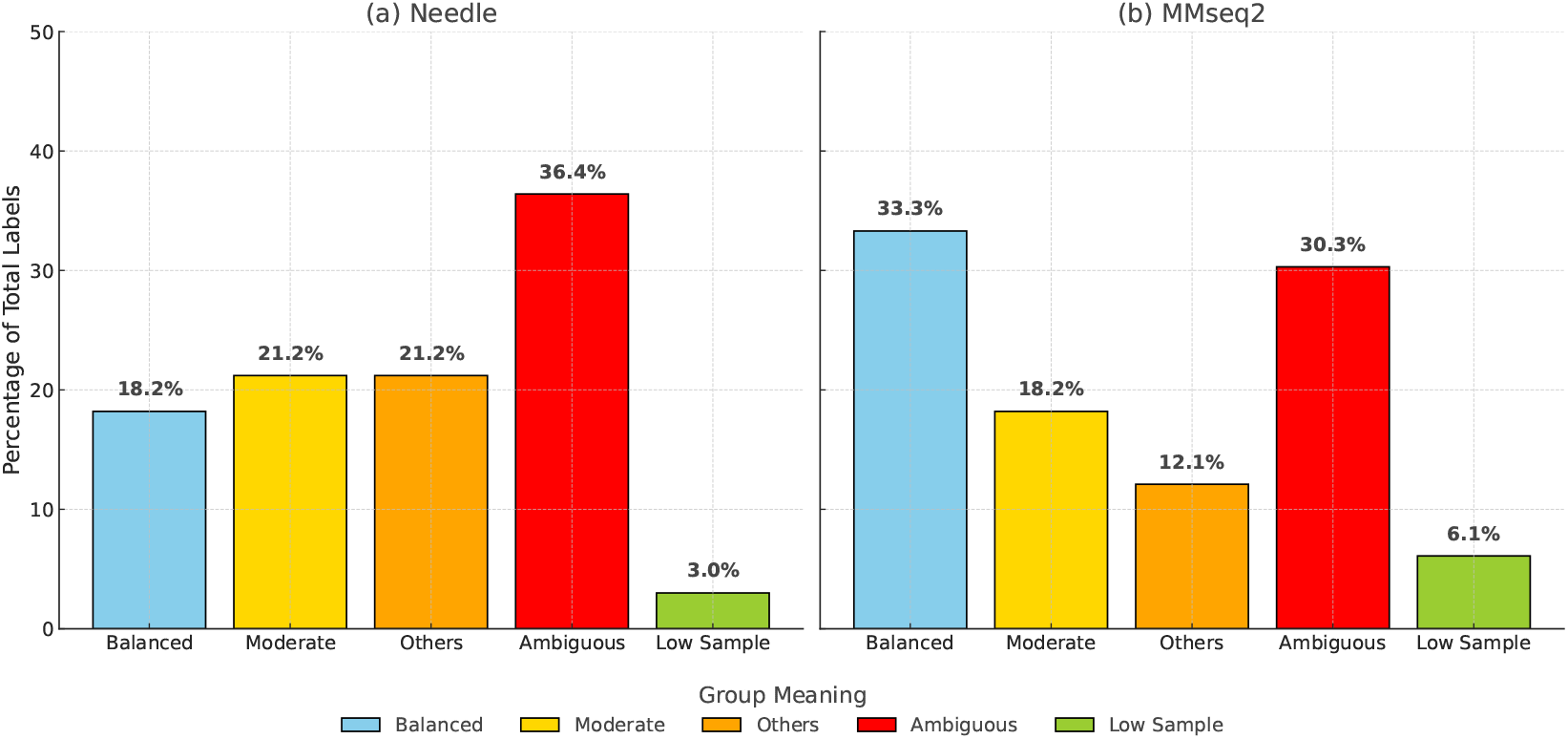
Stratification ratio for GraphPart Needle and MMseqs2 at 30 percent similarity threshold for ARG dataset.

**Fig. 9:**
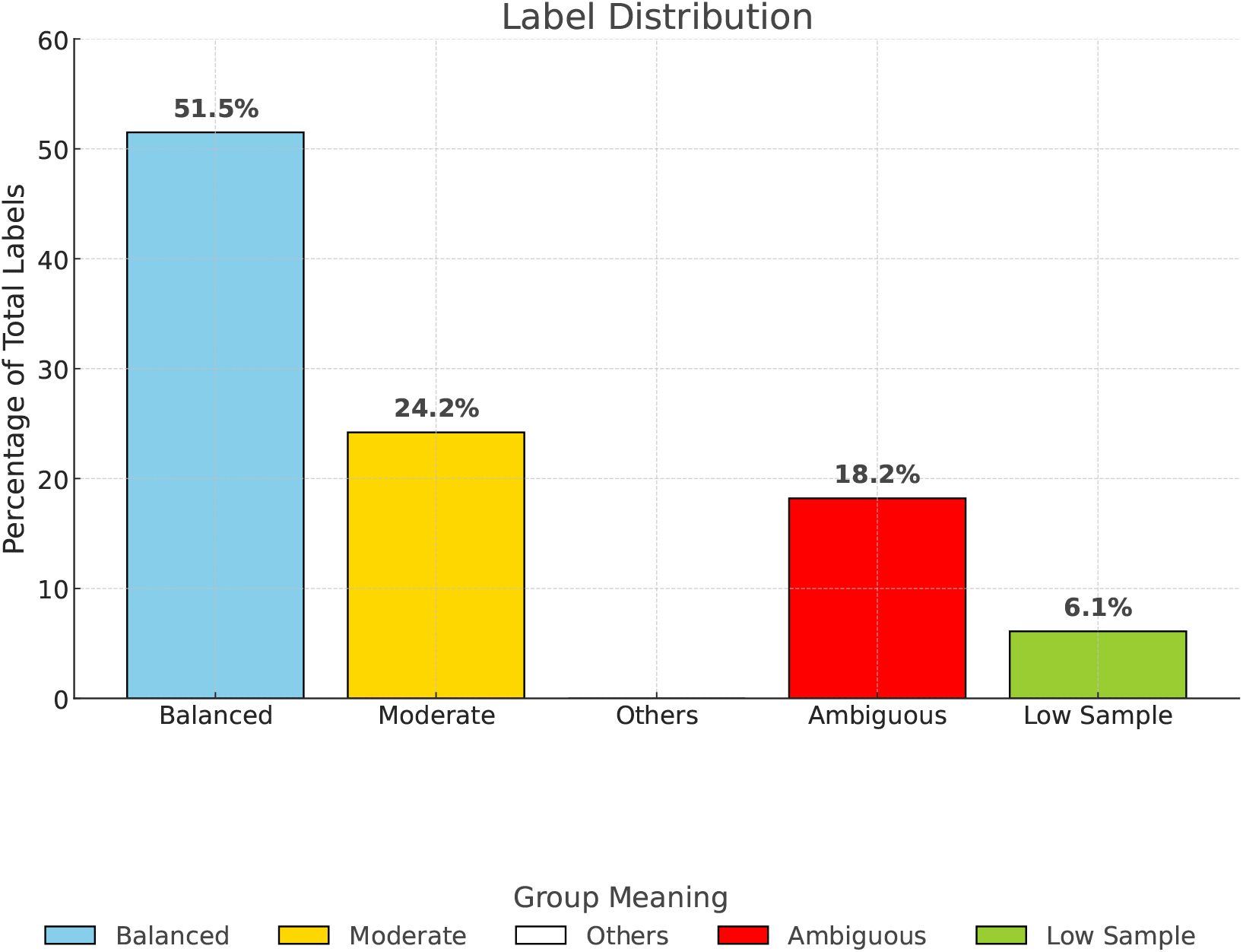
Stratification ratio for Fast-Part at 30 percent similarity threshold for ARG dataset.

### 4.7 Runtime Analysis

Moreover, GraphPart was much slower than Fast-Part, almost 80 times slower with the MMseqs2 mode and 450 times slower for the needle version for the ARG dataset (Table 3). It was impossible to run GraphPart with the needle mode for the mobileOG-db database due to its huge size. All the experiments were done on CPU model name : Intel(R) Core(TM) i9-9820X CPU @ 3.30GHz Cores:20. We used single threads for performing Fast-Part and MMseq analysis. For GraphPart we also used their default parameters.

### 4.8 Sequences Retention

However, GraphPart was able to retain more sequences compared to Fast-Part for the stringent similarity thresholds (Table 4). At stringent similarity thresholds (e.g., 30%), more sequences must be discarded, particularly for small or highly homologous classes. Labels with fewer than 15 sequences or all sequences falling into one cluster may become unsplittable and are either excluded or result in ambiguous partitions.

**Table 4:** Number of sequences removed for 30 percent similarity threshold train-test split for the three datasets by the two models from GraphPart and Fast-Part.

| Dataset | Total Seqs | MMseqs2 | needle | Fast-Part |
| --- | --- | --- | --- | --- |
| ARG | 17282 | 37 | 1 | 823 |
| Deeploc | 14004 | 213 | 0 | 519 |
| NetGPI | 3618 | 1 | 1 | 146 |

While Fast-Part preserves high retention overall, the trade-off between homology separation and sequence inclusion is dataset-dependent. For instance: In the mobileOG-db dataset, 13% of sequences were excluded at the 40% threshold. This is due to the large volume of similar mobile element proteins, which often form large, homologous clusters across different labels.

This exclusion may slightly reduce representativeness, particularly for rare mobile gene families if their sequences are homologous to others and cannot be partitioned. However, the retained 87% still cover all six classes and maintain the intended stratification, ensuring the core structure of the dataset remains intact.

Importantly, we note that excluded sequences are logged and made available for review. Users may choose higher thresholds (e.g., 60% or 80%) for more inclusive splits if redundancy is less concerning.

Fast-Part offers fine-grained control over balance. While some data loss is inevitable at stricter thresholds, the tool’s iterative reassignment mechanism ensures that only non-separable or threshold-violating sequences are discarded, preserving as much as possible without compromising on partition integrity.

### 4.9 Homology Separation

The box plot in Figure 10, compares the maximum pairwise sequence identities observed between different data splits for four partitioning methods, relative to a 40% identity cutoff (red dashed line) under a 60% query-coverage requirement. For Fast-Part and Graph-Part, initial clustering was performed at 40% identity and 60% coverage, followed by an 80:20 train–test split of whole clusters; Fast-Part achieves a median cross-partition identity of approximately 32% and Graph-Part approximately 36%, with only a few outliers above the cutoff, demonstrating tight control of cross-split similarity. MMseqs2 also clustered at 40%/60% before an 80:20 cluster-based split—yields a median of roughly 38% but exhibits a much broader interquartile range and numerous high-identity outliers. CD-HIT, clustered at the same thresholds but randomly assigning clusters to partitions, performs worst, with a median of about 65% and whiskers spanning nearly 0–100%. Sequence similarity was measured using DIAMOND with the same 40% identity and 60% coverage settings. Overall, only Fast-Part and Graph-Part reliably enforce the intended 40% identity separation across data partitions.

**Fig. 10:**
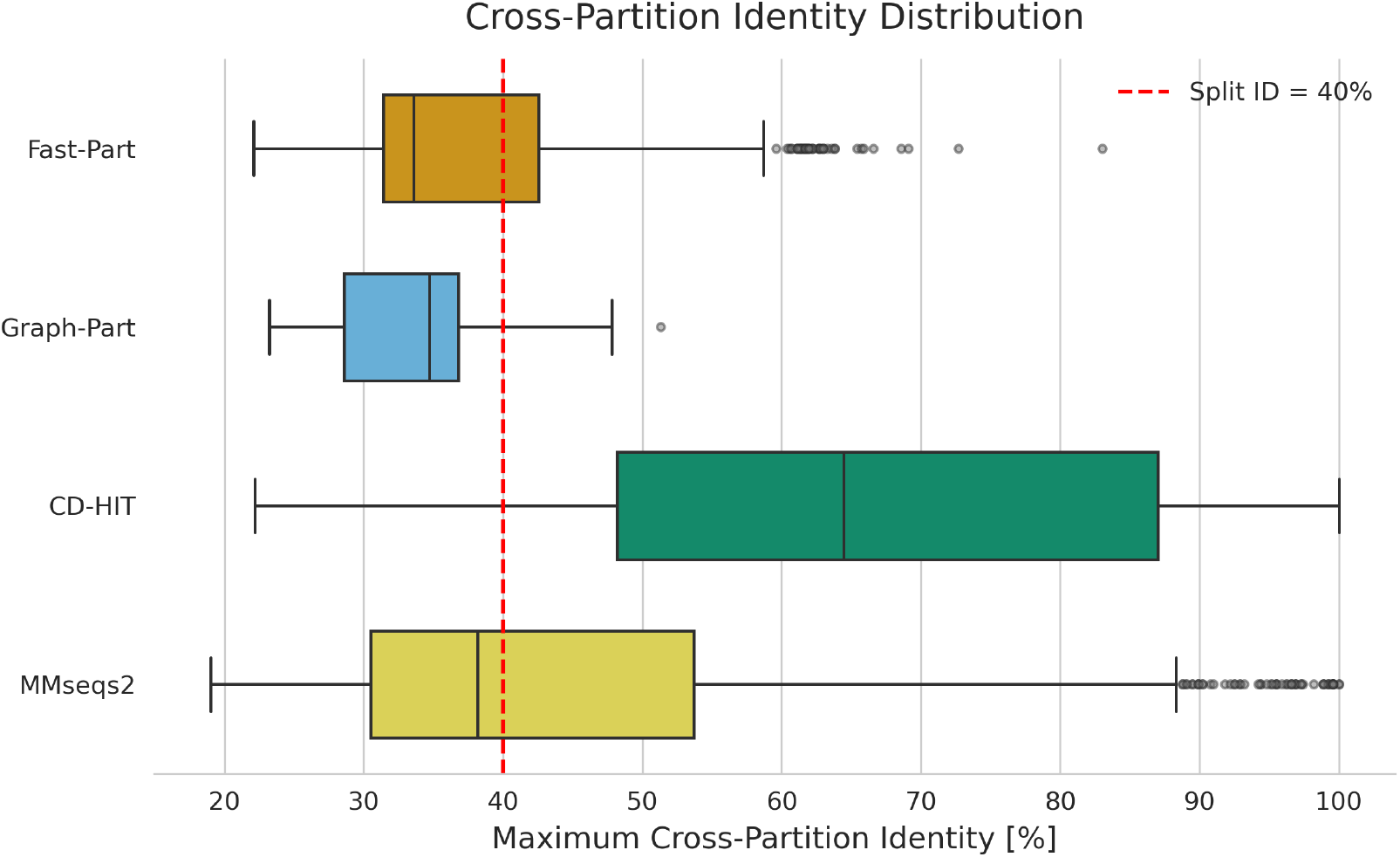
For HMD-ARGDB at 40 percent threshold the homology between cross-partitioned sequences.

For ease of installation, Fast-Part is available both on GitHub and as a Python package via PyPI. The source code and detailed instructions can be found at https://github.com/Shafayat115/Fast-Part. Given a properly formatted FASTA file as input, Fast-Part outputs partitioned training and test files, along with a summary file detailing partition ratios, sequence counts, removed sequences, missing label information, and the configuration settings used for the run. Fast-Part is designed with flexibility in mind, allowing users to input multiple hyperparameters, ensuring it is both powerful and user-friendly. Fast-Part is not applicable for multi-label datasets. While Fast-Part prioritizes data integrity over inclusion, future work may explore merging rare classes with biologically similar categories to preserve label representation. Augmentation techniques to increase representation of low-sample classes for fairer evaluation. Offering an optional relaxed mode that allows slight similarity violations for low-frequency classes under user-defined constraints.

Although Fast-Part was motivated by challenges in ARG classification, its architecture is domain-agnostic and has been evaluated across: DeepLoc (eukaryotic subcellular localization), Metal Resistance Genes (MRGs) (single-label resistance detection), mobileOG-db (large-scale mobile gene datasets). This diversity demonstrates Fast-Part’s applicability to a wide range of biological classification tasks. In principle, any biological task requiring homology-aware train-test partitioning with stratification can benefit from Fast-Part.

## 5 Conclusion

Data partitioning plays a pivotal role in evaluating the performance of sequence classification algorithms, where the accuracy of these methods is heavily influenced by the quality of the dataset splits. Fast-Part offers an advanced solution for partitioning data by ensuring a randomized, yet balanced, distribution of sequences. Fast-Part not only rigorously adheres to the predefined sequence similarity thresholds but also ensures that the distribution of data follows closely the targeted stratification ratios, thus facilitating more reliable and reproducible accuracy assessments in sequence classification tasks.

## Supplementary information

Additional experimental analyses and results have been included in the supplementary materials.

## Acknowledgements

We gratefully acknowledge the support of all funding agencies that made this research possible.

## Declarations

### Funding

This work was partly funded by the National Science Foundation (NSF grants #2319522, #2125798, and #2004751).

### Conflict of interest

The authors declare no conflict of interest.

### Ethics, Consent to Participate, and Consent to Publish declarations

Not applicable.

### Code and Data availability

The source code and detailed instructions can be found at https://github.com/Shafayat115/Fast-Part.

### Author contribution

Shafayat and Muhit contributed to the design of the initial pipeline. Shafayat and Moumi implemented the code and conducted the experiments. Shafayat prepared the manuscript. Liqing supervised the project, reviewed the results, and provided guidance on the manuscript.

## Notes

https://drive.google.com/file/d/1MS6vvMTtMdoG9M2Hi_quMuk9TFEa2iJ-/view?usp=drive_link

### Competing Interest Statement

The authors have declared no competing interest.

### Summary of Updates

New Results, detailed explanation and interpretation.

